# baseLess: lightweight detection of sequences in raw MinION data

**DOI:** 10.1101/2022.07.10.499286

**Authors:** Ben Noordijk, Reindert Nijland, Victor J. Carrion, Jos M. Raaijmakers, Dick de Ridder, Carlos de Lannoy

## Abstract

With its candybar form factor and low initial investment cost, the MinION brought affordable portable nucleic acid analysis within reach. However, translating the electrical signal it outputs into a sequence of bases still requires high-end computer hardware, which remains a caveat when aiming for deployment of many devices at once or usage in remote areas. For applications focusing on detection of a target sequence, such as infectious disease or GMO monitoring, the computational cost of analysis may be reduced by directly detecting the target sequence in the electrical signal instead. Here we present baseLess, a computational tool that enables such target-detection-only analysis. BaseLess makes use of an array of small neural networks, each of which efficiently detects a fixed-size subsequence of the target sequence directly from the electrical signal. We show that baseLess can accurately determine the identity of reads between three closely related fish species and can classify sequences in mixtures of twenty bacterial species, on an inexpensive single-board computer.

**Availability:** baseLess and all code used in data preparation and validation is available on Github at https://github.com/cvdelannoy/baseLess, under an MIT license. Used validation data and scripts can be found at https://doi.org/10.4121/20261392, under an MIT license.

## 1 Introduction

Nucleic acid (NA) sequencing is no longer the costly endeavor it once was; while two decades ago analysis of a single genome could occupy multiple labs over several years [1], technological innovations have now driven the per-base cost down sufficiently to allow routine sequencing for other purposes than scientific discovery, including forensics [2] and clinical diagnoses [3–5]. The case for such usage was strengthened further with the introduction of Oxford Nanopore Technology (ONT)’s MinION, a low-cost, small-size sequencing device. No longer inhibited by high initial investment costs or poor portability, small laboratories and individual users may now opt for in-house sequencing and on-site analysis in remote locations [6–8].

This development was possible due to the introduction of a new sequencing mechanism; rather than the fluorescence-based sequencing-by-synthesis approach employed by previous devices, the MinION sequences DNA strands of arbitrary length by ratcheting them through a nanopore while reading out the electric current [9]. This readout is colloquially referred to as a “squiggle”. As the nucleotide combination residing in the nanopore at a given moment influences the electrical resistance, the squiggle carries information on the sequence. In a process termed “basecalling”, the NA sequence is deduced from the squiggle.

Although the MinION itself is an inexpensive NA sequencer, real-time data analysis currently still requires at least a high-end laptop. For some applications, e.g. the distribution of thousands of devices for infectious disease screening, this may bring along prohibitively high additional costs. It would therefore be beneficial if inexpensive computing hardware could be used instead. Depending on the intended purpose, a computationally lighter analysis pipeline may be a solution. As fast computing hardware is mainly required for basecalling, some basecallers have been developed that trade off lower resource requirements against a decreased basecalling accuracy. DeepNano-blitz [10] is the most recent open-source example of such an implementation, while ONT’s proprietary basecaller guppy has a “fast” running mode for this purpose.

Not all applications require information on the full read sequence however. If only detection of a set of known sequences is required, these sequences could be detected directly in the squiggle instead, potentially reducing the computational load even further. Several direct-from-squiggle sequence detection methods have been proposed. Kovaka *et al*. developed UNCALLED [11], which assigns a probability for each 5-mer potentially matching to each squiggle segment and then compares probable series of 5-mers to a pre-indexed genome to quickly map the read to its likely location. Its original purpose is to facilitate “adaptive sampling”, that is, to rapidly detect the likely origin of a read while the strand is still being sequenced, so that sequencing of strands from non-target sources may be terminated early [12]. UNCALLED can easily be repurposed to perform general sequence detection; however, the index-based approach carries several disadvantages. Efficiency decreases for larger and more repetitive genomes and re-indexing is required to attune the tool to a new target sequence. Moreover, accuracy was found to be low for short sequences [13]. Similarly to UNCALLED, SquiggleNet was designed for adaptive sampling [13]. Following a more straightforward approach, it uses a neural network trained for the recognition of a given genome to decide whether squiggles belong to a species or not. Previously SquiggleNet was found to outperform UNCALLED in terms of both accuracy and processing speed, but the required re-training of SquiggleNet for a given species is a highly resource- and time-consuming process.

Here we introduce baseLess, a computationally efficient and flexible approach to direct sequence detection (Figure 1). Using an array of small neural networks, each pre-trained to recognize a single *k*-mer, baseLess can determine whether a read can be mapped to a given sequence or not. Configuring our tool to detect a sequence requires only the selection of target *k*-mers and their associated pre-trained neural networks. We show that baseLess can perform species detection on eukaryotic whole genome sequencing data against a background of similar species, as well as 16S-based species detection of prokaryotes agnostic of background sequences. BaseLess is more accurate than direct sequence detection pipelines and less computationally demanding than basecalling-and-mapping, allowing it to run on more affordable (∼$100) analysis hardware. As such, it removes an important economical bottleneck for highly distributed and remote field analysis using the MinION.

**Figure 1:**
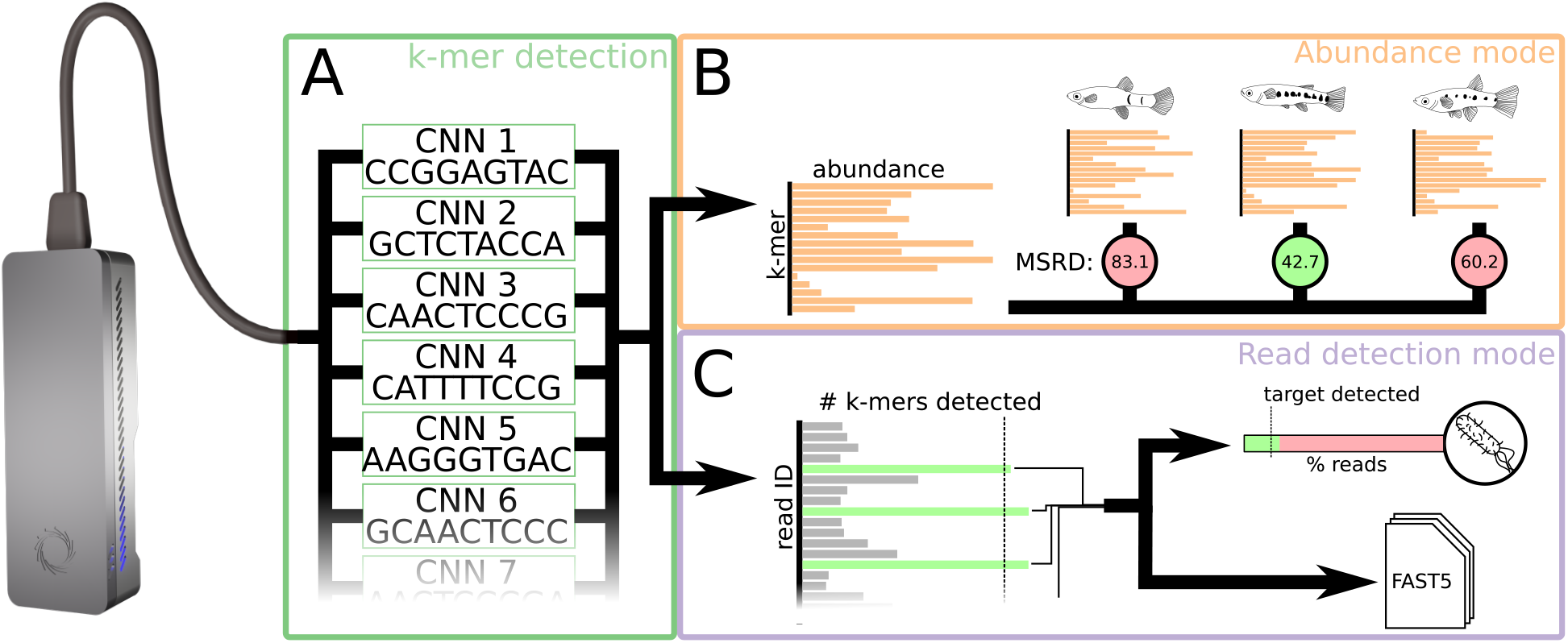
Schematic overview of the baseLess sequence detection tool. **(A)** baseLess detects sequences using an array of pre-trained interchangeable neural networks, each of which detects a specific *k*-mer. These *k*-mers have been specifically selected to allow discrimination of a target sequence. **(B)** In abundance mode, network outputs are summed over all reads and presented as an estimate of *k*-mer abundance. This estimate is compared against genome-based estimates for several closely related species by calculating the mean squared rank difference (MSRD). The species for which the MSRD is lowest is the most likely source of the reads. **(C)** In read-based detection mode, a target sequence is sought in each individual read. A minimum fraction of *k*-mers needs to be detected in a read before it is classified as a target read. A target species is detected if a minimum fraction of analysed reads can be ascribed to it. Reads ascribed to the target species are also stored in a FAST5 file for further analysis.

## 2 Results

### 2.1 Tool structure

BaseLess deduces the presence of a target sequence by detecting squiggle segments corresponding to salient short sequences, *k*-mers, using an array of convolutional neural networks (CNNs) (Figure 1A). Each CNN detects a single *k*-mer, a relatively simple task, thus the network complexity can be kept low. This divide-and-conquer strategy has several advantages. All CNNs can process a read in parallel, which makes baseLess computationally efficient. Furthermore, given a library of pre-trained CNNs, baseLess can easily be reconfigured to detect a different target sequence by combining a different set of CNNs. Finally, sufficient data to train the CNNs is usually available; shorter sequences generally occur more often than longer sequences, thus a read set of any source, once corrected for basecalling errors (see Methods), provides sufficient data to train for a wide range of *k*-mers.

To complete the baseLess network, the outputs of the CNN array are combined using one of two aggregation rules. If configured in “abundance mode”, baseLess returns the number of occurrences found for each *k*-mer, which may then be compared to abundance estimates derived from a target genome (Figure 1B). In “read detection mode”, the network is configured to decide whether a sufficiently large fraction of its *k*-mers have been found in a given read to conclude that it contained the target sequence (Figure 1C). These modes are explained and evaluated in more detail below.

### 2.2 Abundance-based species detection

As baseLess provides fast and accurate inference on low-cost hardware, it is highly suited to determine the species or strain to which a given individual belongs at remote sampling locations, or at many locations simultaneously. Practical applications of such usage may be found in ecological monitoring of visually similar species or forensic investigation of patented crops. For such tasks, baseLess should be configured in abundance mode, which requires the target species’ genome and a set of background genomes – genomes of species from which the target species must be discerned. The *k*-mer set used for discrimination is then found by combining *k*-mers that are highly abundant in the target genome yet found less than average in the background genomes, or vice versa. To determine the origin of a sample, baseLess ranks *k*-mers by abundance as measured in the reads and compares it to their abundance ranking in the target and background genomes, using the mean squared rank difference (MSRD):

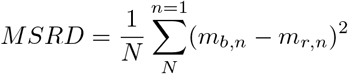

Here *m*_*b,n*_ and *m*_*r,n*_ are the rank for *k*-mer *m* based on abundances in analyzed reads and in a reference respectively. *N* is the total number of *k*-mers analyzed in the reads.

To test baseLess’ performance in this scenario, we analysed unamplified whole-genome MinION reads from three related guppy species: *Phalloptychus januarius, Poeciliopsis gracilis* and *Poeciliopsis turneri* [14]. In three separate analyses, we configured our tool for detection of one of the species against the other two, using Illumina short-read assemblies of the same individuals as target and background genomes to avoid the risk of detecting species based on MinION-specific sequencing errors. We then analyzed a set of 2,000 MinION reads originating from the target species. We found that baseLess consistently calls the correct species for each analyzed readset (Figure 2A-C). Moreover, baseLess did not need the full 2,000 reads for any classification; stable MSRD values were attained after 52, 352 and 84 reads for *P. gracilis, P. januarius* and *P. turneri* respectively. To test whether baseLess indeed detects differences between species and not between individuals, we also ran classification on samples of a family of four *P. gracilis* individuals, using a *k*-mer set selected using the genome of an unrelated *P. gracilis* individual. baseLess consistently called the correct species while requiring less than a hundred reads.

**Figure 2:**
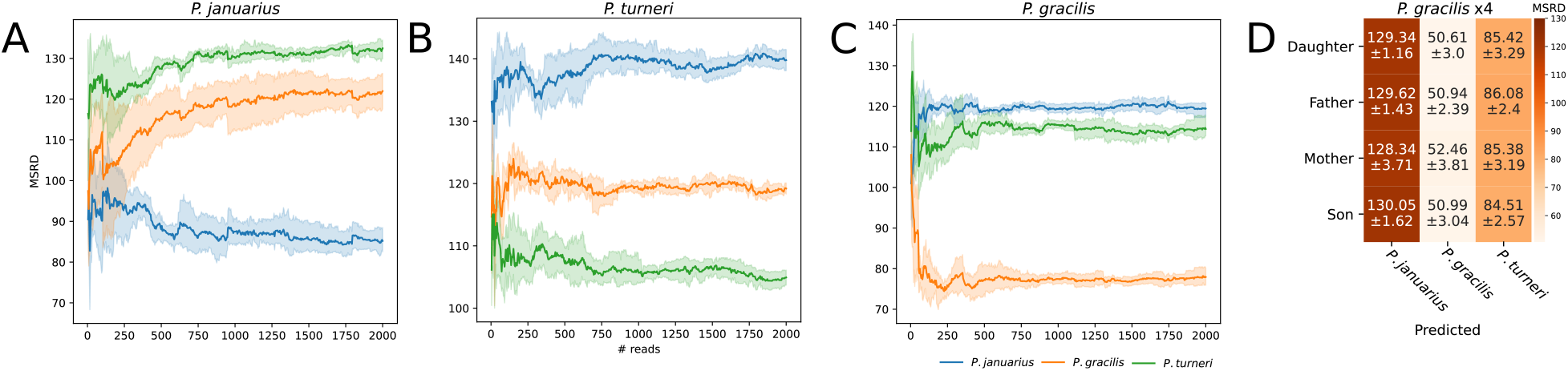
Mean squared rank differences (MSRD) based on a comparison of *k*-mer abundances estimated from reads by baseLess and the abundances in genomes of three closely related fish species. A low MSRD indicates that *k*-kmer abundances in sample and genome are alike and that reads are thus more likely derived from that genome. Results are presented for **(A)** *Phalloptychus januarius*, **(B)** *Poeciliopsis turneri*, **(C)** *Poeciliopsis gracilis* and **(D)** a family of four *P. gracilis* individuals of which no assembled genome was used in the configuration of baseLess. In A, B and C colored areas denote the 95% confidence interval. In D, numbers are formatted as mean *±* standard deviation over 2000 reads.

Interestingly, the *k*-mer rankings also followed the phylogenetic relation between the species; in all detection experiments, MSRD values for *P. gracilis* and *P. turneri* were consistently closer to each other than to *P. januarius*, which is indeed of a different genus. This implies that, even if the genome of the correct species is not included, the relative identity of a sample may be inferred by comparing measured abundances to several related species.

### 2.3 Read-based species detection

In specific applications, a sample may contain a mixture of DNA of many species, from which a species of interest must be detected. Possible scenarios include the screening for infectious disease agents at events or at national borders, or detection of indicator species for environmental health. In 16S cDNA samples, baseLess may be configured to detect such a species of interest by selecting a combination of *k*-mers unique to the target’s 16S sequence, and running it in read detection mode. In this configuration, baseLess detects each *k*-mer on a per-read basis, rather than summing occurrences over all reads as is done in abundance mode. If a minimum fraction of target *k*-mers is found in a read, it is attributed to the target species. The raw squiggle of found target sequences is stored to allow more in-depth analysis at a later stage, while non-target reads are discarded to decrease data storage footprint. To allow reliable detection of a wide range of species against an arbitrary genomic background, we composed a list of *k*-mers which both varied in sequence composition and produced easily differentiable squiggle segments. This list was further filtered to only contain *k*-mers that are present in NCBI 16S sequences, yet sufficiently rare to allow for species discrimination (see Methods).

To test this approach we amplified and sequenced the 16S rRNA regions of an artificial microbial community of twenty-one known species on the MinION (Supplementary table S1). 400,000 reads were fully basecalled and mapped to the twenty-one genomes of the species to determine their likely origin. No reads were mapped to the *Porphyromonas gingivalis* genome, thus this species was left out of subsequent analysis. Read numbers for other species varied between 11 and 51,040.

We reconfigured baseLess and ran inference for each of the species in a five-fold cross validation scheme, to determine how well it could identify the origin of reads. Running speeds were benchmarked on two different classes of hardware; the Nvidia Jetson Nano (2GB), a ∼$100 single-board computer with dedicated GPU (Nvidia Maxwell, 128 cores@921MHz), and a mid-tier desktop computer with dedicated GPU (Nvidia GeForce RTX 3070, 5888 cores@173GHz). To allow straightforward comparison, all tools were given access to 3 CPU cores and the GPU if required. We compared the performance of baseLess on 16S read classification to that of four other pipelines: full basecalling by either DeepNano-blitz [10] or guppy in “fast” mode, followed by mapping using minimap2 [15] (“DeepNano+minimap2” and “guppy+minimap2” respectively); UNCALLED [11]; and SquiggleNet [13].

BaseLess consistently identified its target reads with more than 95% accuracy and an *F*_1_ score of 0.54 on average (Figure 3A), with the exception of *Helicobacter pylori*. Guppy+minimap2 outperformed other pipelines consistently, however compared to DeepNano+minimap2 and UNCALLED, baseLess yielded a higher accuracy and a higher *F*_1_-score for all species. Under default settings SquiggleNet only had sufficient data to classify reads of the three species for which the most reads were available: *E. coli, H. pylori* and *L. monocytogenes*. On these species, baseLess preformed similarly to, or better than Squiggle Net.

**Figure 3:**
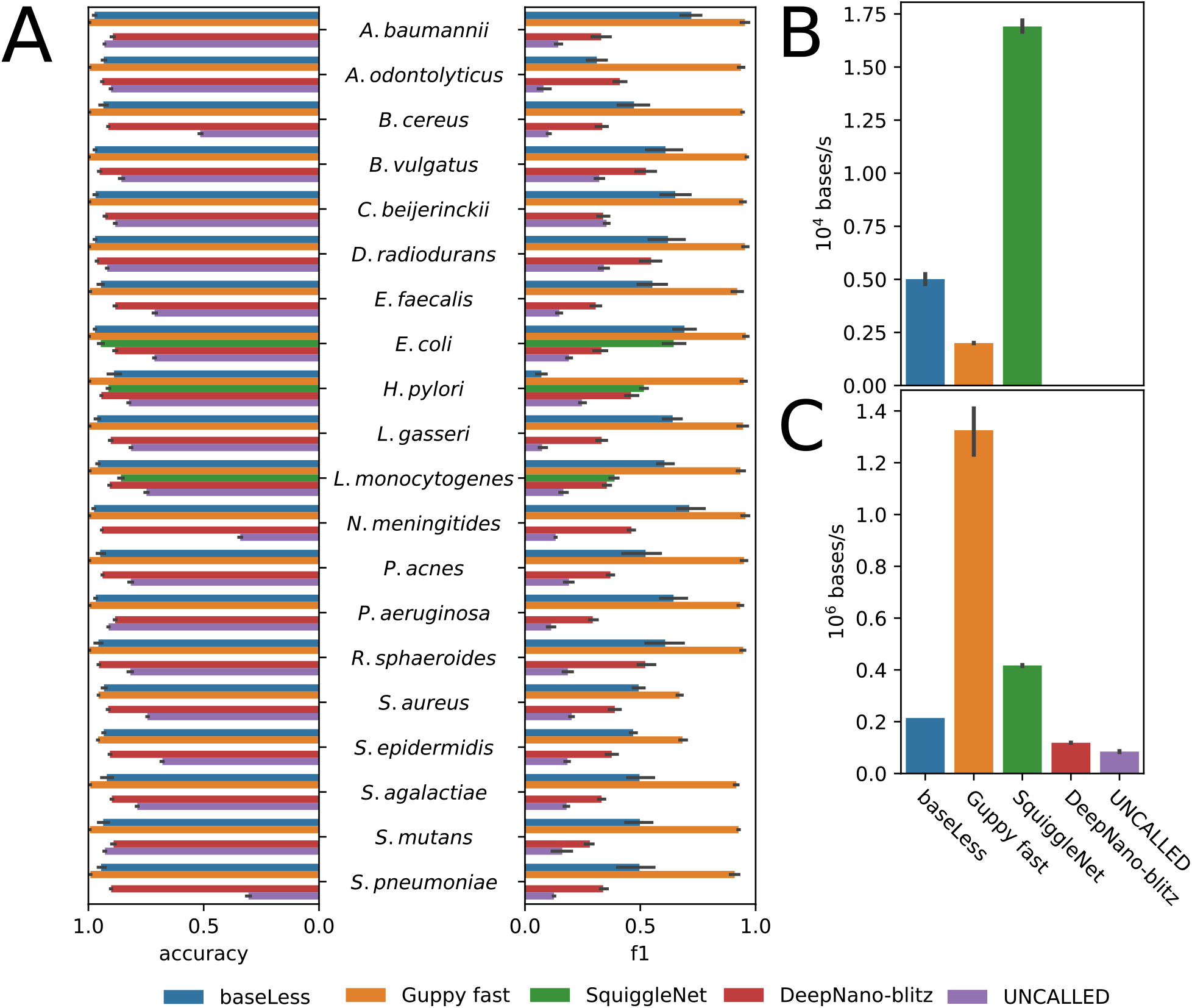
Performance in 16S-based species detection for a community of 20 species, of baseLess and four competing analysis tools; DeepNano-blitz; Guppy (fast mode); UNCALLED and SquiggleNet. As DeepNano-blitz and Guppy are basecallers, not read mapping tools, minimap2 is used to obtain the mapping. **(A)** Accuracy and *F*_1_ score per species for all four tools. Species are sorted by read abundance in the dataset, from high (top) to bottom (low). Black bars denote standard deviation over five cross-validation folds. **(B)** Analysis speed for each tool on the Nvidia Jetson Nano (2GB) single-board computer and **(C)** on a high-end desktop computer. Black bars denote standard deviation over ten analysis runs on 1000 reads.

In the speed benchmark baseLess processed 5.0 kilobases per second (kbps) on the Jetson Nano. We found that baseLess was more than twice as fast as Guppy+minimap2 (2.0 kbps), which could not make use of the Jetson GPU due to software compatibility issues. SquiggleNet ran the fastest at 17 kbps. None of the tools were able to match the theoretical maximum throughput of the MinION (230 kbps). We were unable to install DeepNano-blitz and UNCALLED on this hardware, possibly due to incompatibility with the energy-efficient AARCH64 CPU architecture used in the Jetson Nano and most other single-board computers. As expected, processing speeds were much higher on high-end desktop hardware with the three GPU-accelerated tools – baseLess, guppy+minimap2 and SquiggleNet – performing best. At 1.3 megabase per second (Mbps), guppy+minimap2 was faster than all other tools. SquiggleNet (420 kbps) was again faster than baseLess (210 kbps), which in turn out-competed DeepNano+minimap2 (120 kbps) and UNCALLED (84 kbps).

## 3 Discussion

In this work we proposed a method to identify whole genomes or amplified sequences in nanopore reads by detecting salient *k*-mers using an array of individual, interchangeable neural networks. We show that baseLess, our implementation of this method, is capable of correctly classifying single-species whole-genome sequencing samples, given the target species’ genome and a set of off-target genomes. This is useful for species determination of larger organisms, though not for environmental samples of microbes, which contain many species of which most may be unknown. We therefore also implemented an alternative running mode, which allows microbial species detection against an unknown background, suitable for smaller genomes or PCR-amplified samples.

The world-wide demand for microbe screening, most prominently for infectious disease agents, is currently filled mostly by lateral-flow antibody and qPCR tests. Antibody tests have a turnaround time of minutes, require little training to use and can be mass-produced at low expense, but require a redesign and subsequent re-distribution for the detection of different targets. Moreover, detection is not as reliable as that of nucleic acid analysis [16]. qPCR is generally more reliable, but also requires newly designed primers for different targets. We propose that MinION-based sequence detection using baseLess could fulfill a role similar to that of qPCR; while quick and inexpensive mass produced antibody tests are difficult to improve on for sustained monitoring of a single biological agent, the fast turnaround times and adaptable nature of MinION-based sequence detection would provide an improvement over the logistics and organization required for qPCR testing sites. Reconfiguring baseLess for the detection of a new agent or variant only requires loading the networks for a different set of *k*-mers. Moreover, as found target reads are stored, these may be analyzed in depth afterwards, giving researchers an unprecedented wealth of information on mutations from each detected occurrence of the agent. Especially relevant in this context, though left unexplored here, would be microbe detection using direct DNA or RNA sequencing, as omission of PCR steps would bring down turnaround time even further.

To our knowledge, baseLess is the first tool built specifically to perform MinION-based sequence detection, but other tools can be repurposed to perform the same task. We thus compared baseLess to two fast full basecalling-and-mapping pipelines and two adaptive sequencing tools. Of these competitors, only the guppy+minimap2 pipeline could consistently classify reads with a higher accuracy than baseLess. However, guppy was not optimized for use with low-powered hardware such as the Nvidia Jetson Nano featured in this work, thus it ran slower. This may change if current software compatibility issues are resolved in future releases, although then still the 2GB memory limit of this hardware may prove problematic. SquiggleNet performed similarly to baseLess in terms of accuracy and was more than three times faster on the Jetson Nano. However, due to its high training data requirements it could only be evaluated on three of the twenty species tested here. BaseLess did not suffer from this disadvantage as it only needs examples of *k*-mers to train, which may be obtained from any source. Furthermore, SquiggleNet requires retraining to detect new species, while baseLess only needs reconfiguration for a different set of *k*-mers. We thus argue that baseLess has more potential to be developed into an accurate yet flexible sequence detection tool than its competitors.

Several venues may be explored to further optimize our workflow. Importantly, baseLess’ computational efficiency can be further increased; we ran our tool using Tensorflow, a fully equipped deep learning library, however to run inference on low-powered hardware more efficiently, light-weight frameworks such as Tensorflow-lite and TensorRT may be employed. Further optimization would allow baseLess to analyse reads at a speed more similar to the MinION’s throughput, or preserve computational resources for other tasks, such as driving the MinION itself. Furthermore, we note that the amplification of 16S sequences used in our 16S performance evaluation remains a bottleneck in sequence detection. Instead, the MinION may also be used to directly sequence RNA. As ribosomal RNA makes up a large part of the total RNA content of prokaryotes [17], it would be interesting to evaluate classification based on unamplified RNA content instead.

In summary, the results obtained inspire confidence that using baseLess, the MinION can be turned into a mobile species detector for under $1,000, thus paving the way for large-scale nucleic acid-based detection of biological agents in any environment.

## 4 Methods

### 4.1 Network design procedure

Individual *k*-mers are recognized using 1D convolutional neural networks implemented in Tensorflow 2.3 [18]. We optimized hyperparameters through 100 rounds of training and evaluation on 33, 549 and 3, 241 held-out training and test reads respectively to obtain the final network architecture (Supplementary figure S1). After each round of training and evaluation, the next hyperparameter set was selected using a tree-structured Parzen estimator implemented in hyperopt [19]. The objective function was designed to increase the *F*_1_ score while decreasing network size:

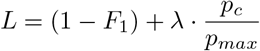

Here *L* denotes the loss to be minimized, *p*_*c*_ denotes the number of parameters in the current iteration of the network and *p*_*max*_ denotes the maximum number of parameters attainable given the boundaries of the parameter search space. The parameter *λ* controls the trade-off between accuracy and network size and was set to 0.01.

Networks output the posterior probability of their target *k*-mers being present in a squiggle segment. The threshold above which this posterior probability is considered sufficiently high to detect the presence of a *k*-mer was chosen to maximize the *F*_1_-score, using a grid search on training data for probabilities between 0.75 and 0.999 with a step size of 0.001. For read detection mode, the fraction of *k*-mers to be detected before the target sequence is considered present must be set as well. This parameter was optimized simultaneously with the posterior probability threshold.

### 4.2 False positive rate simulation

An optimal choice for the value of *k* should balance the abundance of a *k*-mer, such that it is rare enough to discriminate sequences, yet not so rare that it never occurs at all. We approximate this optimal value by considering the probability of detecting target sequences in random sequences by chance.

Assuming all canonical *k*-mers are equally represented, the expected number of *k*-mer occurrences in a read of length *L*_*read*_ is *L*_*read*_ · 2 · 4^−*k*^ and the probability of a *k*-mer occurring in the sequence at least once can be estimated using a Poisson distribution. Assuming we draw networks detecting *k*-mers from a library of pre-generated networks *A*, the expected number of *k*-mers found in a target sequence at least once can be calculated:

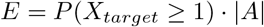

Here *X*_*target*_ is the number of occurrences of a *k*-mer in the target sequence, |*A*| is the size of the *k*-mer network library and *E* is the expected number of *k*-mers in *A* found in the target sequence. A false positive occurs when a read that does not contain the target sequence contains all the selected *k*-kmers of the target read by chance. The rate at which this occurs can be estimated as follows:

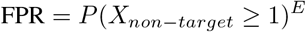

Here *X*_*non*−*target*_ denotes the number of occurrences of a *k*-mer in the non-target sequence and FPR denotes the false positive rate. We performed FPR simulations for different values of *k*, representative values for non-target sequence lengths – 30 and 50 kb, representing full nanopore read lengths – and target sequence lengths – 0.6, 1.5 and 30 kb, representing BOLD barcodes [20], 16S sequences and whole coronavirus genomes respectively – and selected the value for *k* that minimized FPR. Both *k* = 8 and *k* = 9 returned good FPR values, thus we included *k*-mers of both sizes in subsequent steps.

### 4.3 *k*-mer library design

For 16S sequence detection we composed a library of 1,500 suitable *k*-mers, which should allow detection of a wide range of species. Similar to Doroschak *et al*. [21], we used an evolutionary algorithm to select *k*-mers that are dissimilar in sequence and produce easily distinguishable squiggles. To enforce sequence dissimilarity, only *k*-mers with a maximum Smith-Waterman score of 6 (assuming gap penalty, match score and mismatch score of -4, 1 and -1 respectively) to other selected *k*-mers are allowed, while squiggle dissimilarity is enforced by comparing simulated squiggles as produced by guppy (v. 5.0.11+2b6dbff). That is, we only accept modifications made to *k*-mers by the evolutionary algorithm if both the minimum and the average dynamic timewarping score between its squiggle and the other squiggles in the set increase. Furthermore, for the bacterial case study, we remove the outer 10 percentiles of most abundant *k*-mers based on 20,959 16S rRNA sequences obtained from NCBI (Bioproject:PRJNA33175) because these *k*-mers are excessively rare or ubiquitous. Additionally, *k*-mers containing four or more of G/C or 5 or more of A/T in a row are rejected as the length of homopolymer stretches can be difficult to detect in squiggles. Starting with a set of random sequences, we ran the evolutionary algorithm for ten rounds of decreasing numbers of proposed mutations per sequence; the initial two rounds applied 5 mutations in each sequence, after which the number of mutations decreased by one for each two rounds.

### 4.4 Nanopore sequencing

*Poeciliidae* reads were obtained from a previous study and have been obtained as described in [14]. For 16S reads, we sequenced pre-made DNA isolate of microbial mock community A (v3.1, HM-278D, BEI resources) on a MinION (Mk.1B, Oxford Nanopore plc.) using accompanying flowcell (FLO-MIN106) and 16S sequencing kit (SQK-RAB204).

### 4.5 Data preparation

All reads were basecalled using guppy (v5.0.11+2b6dbff) in high-accuracy mode. To obtain a ground truth species assignment for 16S reads, we mapped them using BLASTN (v2.9.0+) to the expected twenty-one bacterial GenBank genomes (Supplementary table S1). The species to which the sequence identity was highest was selected as the ground truth species for that read. To correct sequencing errors and assign individual bases to each squiggle segment, we aligned reads to reference genomes using tombo (v1.5.1). *P. gracilis, P. januarius* and *P. turneri* reads were aligned to genomes constructed from the nanopore reads, while 16S reads were aligned to their respective GenBank genomes. These genomes were also used for salient *k*-mer detection in the evaluation of read detection mode. For abundance mode validation, *k*-mers were selected from GenBank short read genomes, built from Illumina reads of the same three individuals (GCA_903067085.1, GCA_902982915.1, and GCA_903068135.1 for *P. gracilis, P. januarius* and *P. turneri* respectively).

### 4.6 Benchmarking

We compared BaseLess performance on 16S reads to four other tools; UNCALLED (v2.0-127-g0fc1cab), SquiggleNet (v1.0), DeepNano-blitz (v1.0) and Guppy (v5.0.11+2b6dbff, “fast” mode). As the latter two tools are basecallers and not mapping tools, the basecalled reads returned by these were mapped to target genomes using minimap2 (2.17-r941) to produce the final prediction.

We performed accuracy and F1 score benchmarks in stratified 5-fold cross validation on 335,000 reads. Tools were run on a PowerEdge R740 server (Dell), on three Xeon Gold 6242 CPUs @2.80GHz (Intel). As Guppy and SquiggleNet were optimized for GPU usage, they were run on a Tesla T4 GPU (NVIDIA). We ran all tools in a Snakemake [22] workflow.

Speed benchmarks were performed on two systems, an Nvidia Jetson Nano System-on-Module (2GB RAM, ARM CPU, 4 cores@1.43GHz, Nvidia Maxwell GPU, 128 cores@921MHz) and a mid-tier desktop computer (32GB RAM, AMD Ryzen 3700x CPU, 16 cores@3.6GHz, Nvidia GeForce RTX 3070, 5888 cores@173GHz). Inference was performed 10 times per tool over 1,000 reads. Tools were given access to 3 CPU cores and the GPU if they were configured to use it.

## 5 Acknowledgements

We thank Elio Schijlen and Bas te Lintel Hekkert for help with nanopore sequencing. We also thank Henri van Kruistum for the provision of raw nanopore reads and nanopore assemblies for *P. gracilis, P. januarius* and *P. turneri*.

