## Supplementary data for "baseLess: lightweight detection of sequences in raw MinION data"

### BASELESS: LIGHTWEIGHT DETECTION OF SEQUENCES IN RAW MINION DATA
### SUPPLEMENTARY MATERIAL

BIORXIV VERSION

**Ben Noordijk**[1], **Reindert Nijland** [2], **Victor J. Carrion**[3,4], **Jos M. Raaijmakers** [3,4]
**Dick de Ridder**[1] **and Carlos de Lannoy** [1,5]*
[1] Bioinformatics Group, Wageningen University, Wageningen, The Netherlands
[2] Marine Animal Ecology, Wageningen University; Wageningen, the Netherlands
[3] Institute of Biology, Leiden University, Leiden, The Netherlands
[4] Department of Microbial Ecology, Netherlands Institute of Ecology, Wageningen, The Netherlands
[5] Department of Bionanoscience, Delft University of Technology, Delft, The Netherlands

#### 1 Supplementary material

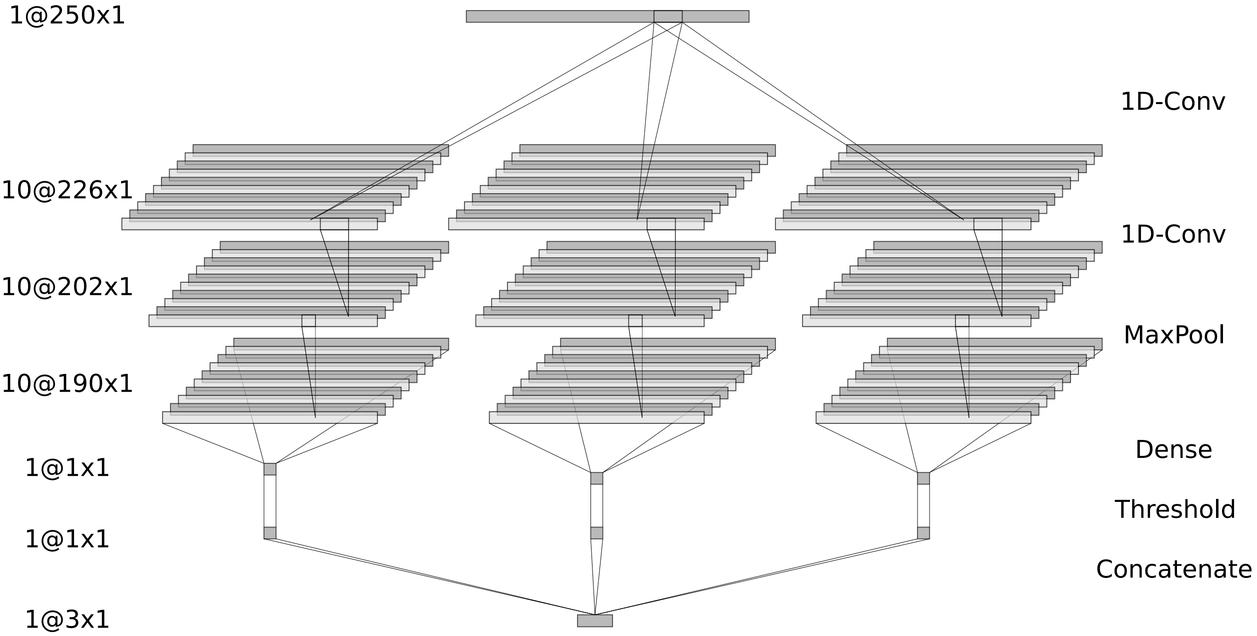


Figure S1: BaseLess neural network structure for the recognition of different $k$-mers. A network for three $k$-mers is shown, although the structure can be expanded to an arbitrary number of $k$-mers. Starting from a squiggle segment, two 1D-convolution layers, a maxpool layer and dense layer are applied consecutively. A threshold on posterior probability is applied to produce a boolean indicating presence or absence of the target $k$-mer, after which resuts are concatenated. In abundance mode concatenated results are summed over all batches (not shown), while in read detection mode an additional rule is added to return a single boolean indicating whether the read contains a predefined minimum fraction of all target $k$-mers.

Table S1: The 21 bacterial strains and associated GenBank assembly accessions used in the validation of the read detection mode of baseLess.

| **Species** | **GenBank Accession** |
|---|---|
| *Acinetobacter baumannii*, strain ATCC 17978 | GCA_013372085.1 |
| *Actinomyces odontolyticus*, strain ATCC 17982 | GCA_000154225.1 |
| *Bacillus cereus*, strain ATCC 10987 | GCA_000008005.1 |
| *Bacteroides vulgatus*, strain ATCC 8482 | GCA_000012825.1 |
| *Clostridium beijerinckii*, strain NCIMB 8052 | GCA_000016965.1 |
| *Deinococcus radiodurans*, strain R1 (smooth) | GCA_000008565.1 |
| *Enterococcus faecalis*, strain OG1RF | GCA_004006275.1 |
| *Escherichia coli*, strain K-12 MG1655 | GCA_000005845.2 |
| *Helicobacter pylori*, strain 26695 | GCA_000008525.1 |
| *Lactobacillus gasseri*, strain ATCC 33323 | GCA_000014425.1 |
| *Listeria monocytogenes*, strain EGDe | GCA_000196035.1 |
| *Neisseria meningitides*, strain MC58 | GCA_000008805.1 |
| *Porphyromonas gingivalis*, strain ATCC 33277 | GCA_000010505.1 |
| *Propionibacterium acnes*, strain KPA171202 | GCA_000008345.1 |
| *Pseudomonas aeruginosa*, strain PAO1-LAC | GCA_000006765.1 |
| *Rhodobacter sphaeroides*, strain ATH 2.4.1 | GCA_000012905.2 |
| *Staphylococcus aureus*, strain USA300_TCH1516 | GCA_000017085.1 |
| *Staphylococcus epidermidis*, strain ATCC 12228 | GCA_000007645.1 |
| *Streptococcus agalactiae*, strain 2603 V/R | GCA_000007265.1 |
| *Streptococcus mutans*, strain UA159 | GCA_000007465.2 |
| *Streptococcus pneumoniae*, strain TIGR4 | GCA_000006885.1 |
